# Reproductive success of inbred strain MV31 of the ctenophore *Mnemiopsis leidyi* in a self-sustaining inland laboratory culture system

**DOI:** 10.1101/2024.07.03.601798

**Authors:** Pranav Garg, Cameron Frey, William E. Browne, Steven S. Plotkin

## Abstract

Ctenophores are an attractive lineage for studying animal evolution due to their early divergence from other metazoans. Among Ctenophora, *Mnemiopsis leidyi* is a model system for developmental, cellular, molecular genetic, and evolutionary studies. Until recently, many of these studies were conducted on wild-caught animals, limiting access to researchers on the coast. Here we present significant advancements towards culturing *M. leidyi* in laboratories without coastal access, enabling its wider use as an experimental and genetic model system. We detail updated feeding regimes that take advantage of co-culturing *Brachionus* rotifers with *Apocyclops* copepods, and quantify the reproductive output of our *M. leidyi* lab strain on this diet. Our updated feeding regime maintains reproductive fitness comparable to wild-caught individuals. Importantly, we have eliminated the logistical complexities and costs of regularly feeding live larval fish to *M. leidyi*. Our updated protocols make it feasible to maintain continuous ctenophore cultures independent of access to both coastal populations of wild *M. leidyi* and larval fish culturing facilities.

## Introduction

Access to a diversity of animal model systems is necessary to understand the origins and evolution of molecular mechanisms that have produced the wide array of specialized cell types unique to metazoans [1]. Ctenophores have been used for over a hundred years to investigate animal biology [2–4]. However, only recently have methods been developed for reliably closing their life cycle in the laboratory [5–7]. The early divergence of the ctenophore lineage from other animals make them a highly informative group for understanding metazoan evolution [8–11].

Ctenophores exhibit a number of interesting biological features, including regeneration [12–15], bioluminescence [16–18], rows of giant cilia used for locomotion [19–21], proprioceptive organs [22–24], immune cells [25–27], functional through-gut [28], prototypical nerve nets [29–31], and adaptations to extreme environments [32]. The genomes of several ctenophore species have now been sequenced and annotated [11, 29, 33–35]. The varied and unusual morphologies of ctenophores have also made them popular for display in public aquariums [36].

The lobate ctenophore, *Mnemiopsis leidyi* [37] (synonymous with *Mnemiopsis mccradyi*), is of particular concern as a highly invasive species capable of significant ecological damage [38, 39]. For example, their invasion of the black sea resulted in the depletion of commercial fish stocks with an economic cost estimated in the billions [40].

The regular seasonal coastal Atlantic availability of *M. leidyi* has enabled research using wild-caught animals in select locations [41]. However, wild-caught animals can introduce uncertainty and variability due to genetic heterogeneity and unknown environmental exposures. A critical but challenging step in developing a model organism is housing and propagation in a controlled environment [42].

Soft bodied planktonic animals that are grown in controlled environments can be kept in planktonkreisel aquariums [43], which maintain a circular flow of water, keeping animals suspended in the water column. A simpler U-shaped “pseudokreisel” [44, 45] design is also popular. Facilities with easy coastal access can use filtered sea-water, while laboratories without coastal access typically use commercially available artificial sea water. The aquaria are fitted with biological and/or chemical filtration equipment to maintain water quality [6, 45, 46].

Several ctenophore species have been sustained in artificial conditions to varying degrees of success [6, 36, 47–49]. Wild caught *M. leidyi* can be held in captivity for weeks and used to obtain embryos and cydippid larvae for developmental studies [7, 50, 51]. Recently, a few methods to continuously culture *M. leidyi* have been published [6, 36, 52]. Maintaining long term cultures requires addressing the challenge of obtaining viable and healthy cydippid larvae from adults living in an artificial habitat. The diet provided to the larvae must be nutritious, as well as appropriately sized for capture and feeding.

Rotifers have been used extensively in aquaculture and zebrafish laboratories as a first food for fish larvae [53, 54]. *Brachionus plicatilis* (“L type”) and *Brachionus rotundiformis* (“S type”) are commonly used for this application. These rotifers can be sustained using live algae and a variety of algae- and yeast-based pastes or dry powders. Ultimately, the nutritional value of rotifers as food depends heavily on their diet [55–57].

Existing protocols have used cultures of *Brachionus* rotifers as the primary source of nutrition for ctenophores. To maintain high fecundity, rotifer based diets have been augmented with larval fish such as Zebrafish [6, 36], or live mysid shrimp [7]. This increases the complexity of maintaining long-term cultures as it requires additional logistical demands for regular sourcing of mysid shrimp and/or larval fish (including adherence to any regulatory requirements associated with vertebrates). While frozen wild-caught copepods have been experimented with [52], they present challenges in maintaining water quality due to the decomposition of uneaten copepods. The culturing protocol we describe here addresses this issue by using live *Apocyclops panamensis* copepods.

Copepods are generally more difficult to grow than rotifers due to their longer development time, with multiple naupliar stages [58]. Many copepod species rely on live algae feeds rather than concentrated algae feeds, which are much more convenient to use [59]. Marine copepods from *Tigriopus, Tisbe, Acartia, Pseudodiaptimus* and *Parvocalanus* genera are popular for use in aquaculture (see e.g. [60] for culturing techniques for several species). In the culturing protocol described here, we chose *Apocyclops panamensis* because they can be sustained on the same algal concentrate used for rotifers. Further, since their temperature requirements are similar to *Brachionus rotundiformis*, we found that they can be stably maintained in a mixed culture with this rotifer indefinitely, so long as maintenance procedures are followed. *Apocyclops royi* is another possible choice due to its much shorter development time [61] among other attractive characteristics including tolerance for high growth densities and synthesis of polyunsaturated fatty acids [62].

In the following, we describe the establishment of a simplified multigenerational inland ctenophore culturing system. We also quantify and compare the reproductive capacity of this established *M. leidyi* laboratory strain, the Miami-Vancouver F31 or MV31 strain, with historical data from wild-type *M. leidyi* available in the literature, as well as against expectations based on theoretical considerations of bioenergetics.

## Results

A laboratory strain of *M. leidyi* originating from a single wild-caught individual from Biscayne Bay Florida in January 2016 was generated through repeated inbreeding [6]. This captive strain has been propagated through self-fertilization, with each subsequent generation reproduced from the self-fertilized spawn of a single individual of the previous generation [6]. A pool of F31 individuals was shipped from Florida to Vancouver, BC in October 2020, and has since been genome-sequenced and maintained in culture over multiple generations as the Miami-Vancouver F31 or MV31 strain as per the protocols presented in the Methods section.

To record the reproductive output and assess the viability of spawned embryos, we measured the sizes of reproductive adults, the number of eggs shed and the number of spawned embryos that hatched from the chorion as presumptively viable larvae. A plot of the number of eggs laid against the adult oral-aboral length (Fig. 1A) shows an increasing trend. The number of hatched larvae *vs*. the spawn size is plotted in Fig. 1B. The plot is well-fit by linear model assuming a zero intercept, with a slope of 0.73 (95 % confidence interval is 0.70–0.77, see Methods). This indicates that approximately 73 % of embryos successfully develop into hatched cydippid larvae under laboratory conditions.

**Figure 1:**
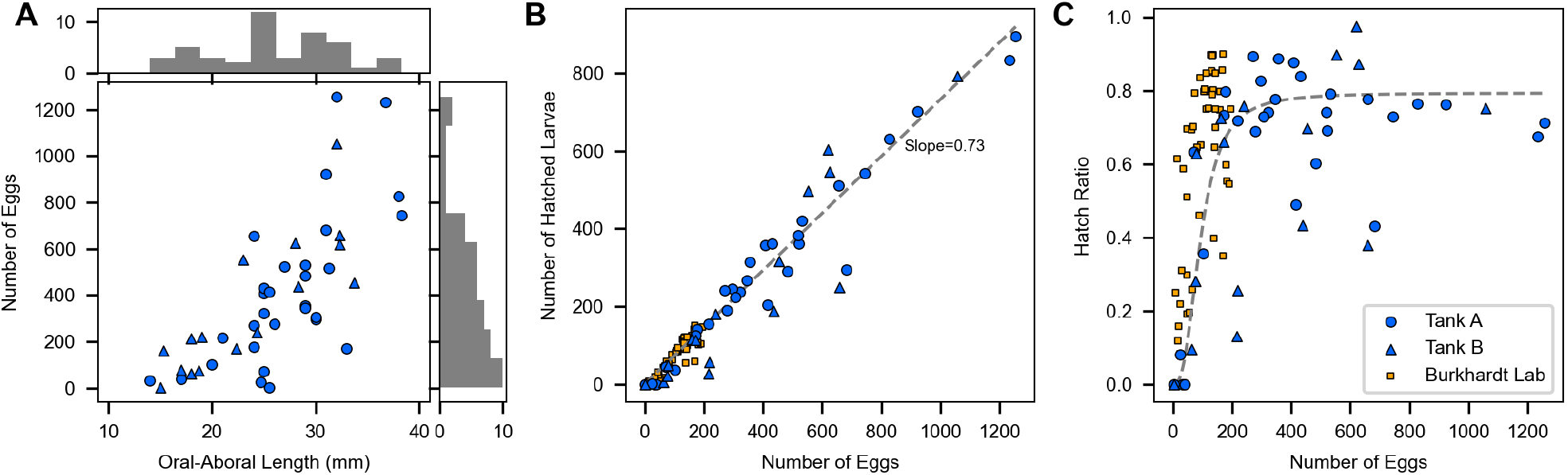
Spawn viability of lab-raised *M. leidyi*. In all panels, animals coming from our pseudokreisel tank “A” are plotted as black circles and those from tank “B” are plotted as blue triangles. Data from [52] is shown as orange squares in the latter two panels. (A) For every adult in our study, the number of eggs spawned is plotted against the oral-aboral length of the animal. (B) The number of hatched larvae is plotted vs. the number of spawned embryos from each animal. A linear fit to the data from our two tanks, assuming zero intercept, yields the function *y* = 0.73*x*, indicating 73% viability. (C) The same data as in panel B, now plotted as the fraction of hatched larvae by dividing by the number of spawned embryos, *vs*. the number of spawned embryos. The light grey curve is a Hill function *y* = *y*_∞_*x*^*n*^/(*a*^*n*^ + *x*^*n*^) plotted as a guide to the eye. A best fit of the function to our data yields the parameters *y*_∞_ = 0.79 (similar to the fit in panel B), *a* = 94, and *n* = 2.7.

The fraction of surviving cydippid larvae may be obtained by dividing the number of larvae that hatched from the chorion by the number of eggs shed. This fraction is plotted *vs*. the number of eggs shed in Fig. 1C. The figure shows a distinct depression in larval viability for small spawn sizes, which is not readily evident in panel (B). We speculate that the nutritional and/or energetic deficits that result in low spawn size are also reflected in the loss of fidelity of embryonic development, perhaps due to improper maternal loading or external environmental factors, resulting in reduced hatching viability. The asymptotic hatch ratio is consistent with the slope in Fig. 1B. The largest values of viability (hatch ratio) are for intermediate spawn sizes in the range of 300-600 embryos, however the decrease in viability with larger hatch sizes is relatively modest. This indicates that any potential reduction in viability due to crowding of embryos in the spawning bowls is a modest effect.

### Comparisons of Reproductive Output with Past Reports

It is interesting to compare the fecundity of this multigenerational lab strain of ctenophores to previous wild caught samples of *M. leidyi* and animals grown for shorter periods in labs. Soto-Àngel et al. [52] present the number of eggs and their viability for laboratory cultured *M. leidyi* individuals. Data from this study is presented alongside our counts in Fig. 1 panels B and C for comparison. These data points were not included in the calculation of best-fit curves. We note that animals grown under the protocols of Soto-Àngel et al. [52] achieved a slightly higher hatch rate than ours, but the overall spawn size was smaller.

While Soto-Àngel et al. did not measure the size of the ctenophores, some previous studies have observed a correlation between the number of spawned eggs and the size of the ctenophore [63–65]. We compared the following four sets of literature data besides our own (Supplementary table S1). Sasson and Ryan [65] have plotted the spawn number *vs*. ctenophore length. Similarly, Baker and Reeve [63] have plotted the spawn number *vs*. length for wild-caught *M. leidyi*, and have also provided tabulated data of both length and egg production for 6 lab-raised *M. leidyi* specimens. Reeve et al. [64] have made direct counts of egg production for two series of growth efficiency experiments at various copepod concentrations. Through the following reasonable modeling assumptions, we were able to convert measurements of ctenophore lengths or dry weights to ctenophore volumes.

We found that our measurements of aboral-lobe length *L*, and lobe-lobe width *D* satisfied the following scaling relationship (Fig. S6A):

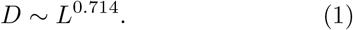

The strong correlation implies *D* may be predicted from *L*, and the sublinear exponent indicates that larger ctenophores become more prolate as they grow, i.e. the length increases more than the width. With this scaling, we expect the volume to scale with length with an exponent of ≈ (2 × 0.714 + 1), i.e.

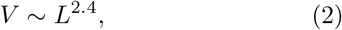

which is borne out by directly plotting the calculated ctenophore volume *V vs*. the length *L* (Fig. S6B), yielding a scaling relationship of

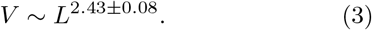

We used this length-volume relationship to predict ctenophore volume from literature data of measured lengths.

To convert between measurements of dry carbon mass *m* and length *L*, we used data from table 3 of Reeve et al. [64] and found that these values satisfy (Fig. S6C)

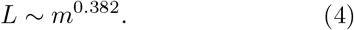

Together with the *V* ∼ *L*^2.43^ scaling between volume and length (Eqs. (3) and (4)) we therefore convert between the measured dry carbon mass and calculated wet volume with a nearly linear relationship, as would be expected:

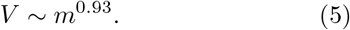

All data are plotted in Fig. 2. We observed an increasing trend between spawn number and ctenophore volume. The data for the set of wild-caught ctenophores from Baker and Reeve [63] deviate from the other data, having higher spawn numbers. This is not unreasonable, as the diet and ecology of wild-caught ctenophores may result in systematically improved spawn numbers. The data from Reeve et al. [64] also deviated noticeably, with individual animals having systematically higher volumes for their respective spawn sizes. This is also unsurprising, given the number of modeling assumptions used to obtain the volume for this dataset. If we remove these two outlying datasets, the remaining three datasets overlap well, and show a scaling relationship between spawn number and volume:

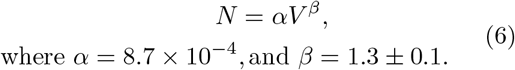

**Figure 2:**
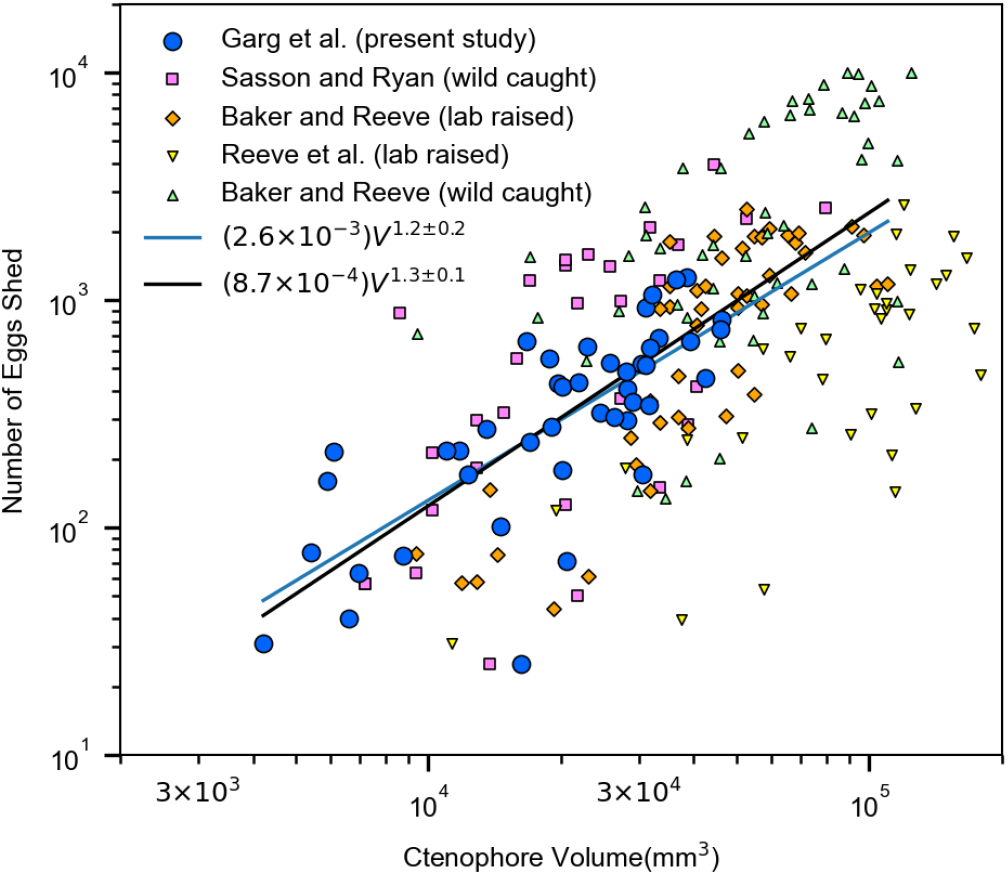
Reproductive output of *M. leidyi*. The number of eggs spawned from each animal is plotted vs. their predicted volume (mm^3^) as obtained from their aboral-lobe lengths (see text). Five datasets from various labs consisting of both wild-caught and lab-raised animals are plotted, including the dataset from the lab strain described here. The linear fit to our dataset exhibits a spawn number scaling with volume as *V* ^1.2*±*0.1^, while a fit to all the data while excluding both the wild caught strain of Baker and Reeve [63] and the lab strain of Reeve and Kremer [64] exhibits a spawn number scaling with volume as *V* ^1.3*±*0.1^ (see text for description).

**Figure 3:**
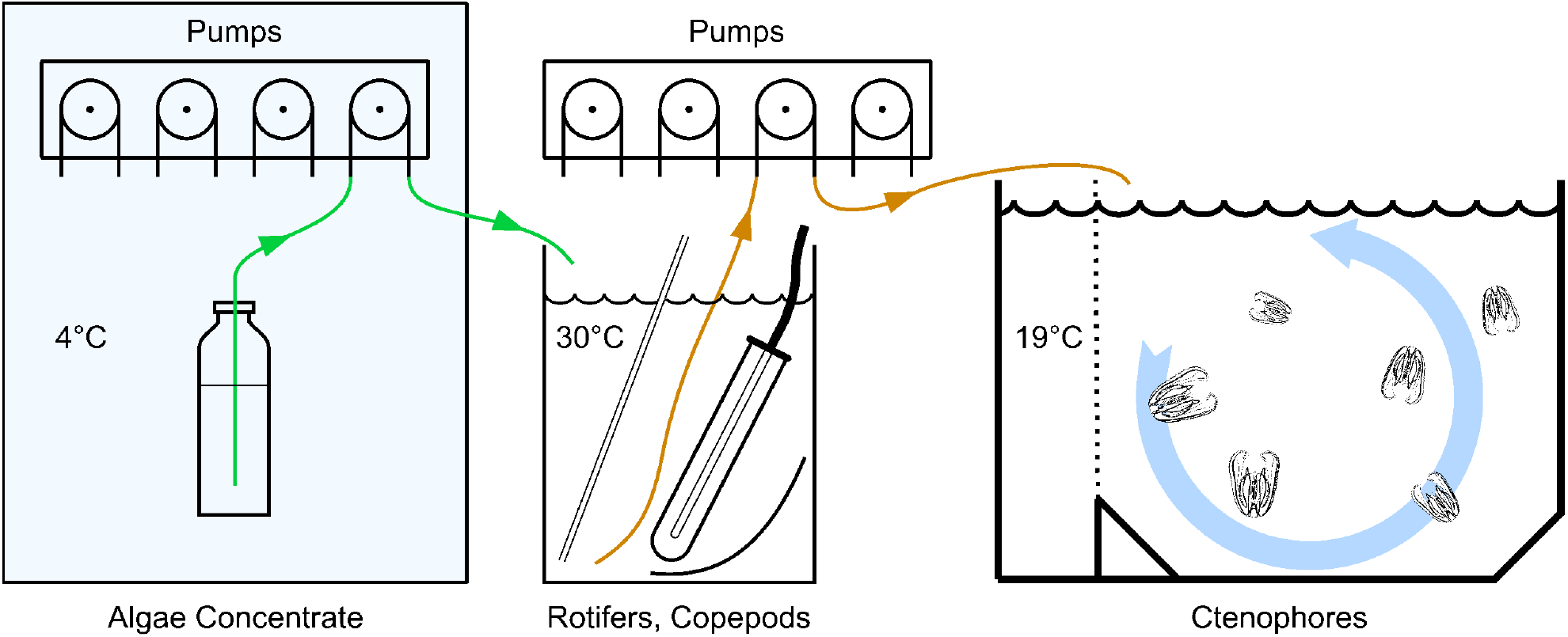
Schematic of the culture system. A chilled algae concentrate is fed to a mixed zooplankton culture of *Brachionus rotundiformis* rotifers and *Apocyclops panamensis*, which in turn is fed to a culture of *M. leidyi*. The algae concentrate may be supplemented with live algae, as described in the text. System components are described in detail in Supplementary section S1.

### Analysis of Metabolic Output

Beyond enabling comparisons across datasets, the above analysis allows us to comment on the metabolic output of *M. leidyi*, and the corresponding metabolic cost of spawning. It is reasonable for the metabolic output of an animal to increase with animal size. In fact, the total metabolic (respiration) rate *R* has been observed to scale with body mass *M* as *R* = *a* ·*M* ^*b*^, with b=3/4 [66]. This allometric scaling law, the so-called “3/4-power law”, has been observed to span across a range of animal phyla and over 16 orders of magnitude of body mass [67, 68]. Theories have been proposed to explain the scaling exponent through the requirements of nutrient distribution and transport throughout an organism [69].

On the other hand, within a given species, the intraspecific scaling exponent of metabolic rate vs body mass often shows significant deviation from the 3/4-power scaling law [70, 71]. The allometric coefficient between metabolism and mass has been measured for *M. leidyi* by Kremer [72] to be *b* = 0.96 ± 0.18 at 20 °C. This is significantly different from both a body-surface area exponent of 2/3, and as well, the 3/4 power scaling law [71], indicating that the relationship between metabolism and mass is distinct from one correlating directly with these physical measurements of size.

We can think of the spawn number *N* as a kind of metabolic output, which may have a different scaling with mass than the respiratory output *R* above. In addition, the volume can be considered a rough proxy for mass, which is somewhat justified given that the scaling relationship between volume and dry mass above was found to be *V* ∼ *m*^0.93^, which is fairly close to linear scaling. With these interpretations in mind, our scaling relationship between spawn number and volume was found to be superlinear, with an exponent of *β* = 1.3 ± 0.1 (Eq. (6)), implying that more resources may be allocated to spawning per volume (or per mass) as ctenophores grow larger. It is worth noting that the scaling law for wild caught ctenophores from Sasson and Ryan [65] is consistent with the above number, having an exponent *β*_wild_ = 1.4 ± 0.3. If we consider only our measured data alone, the best fit becomes *N* = *α*^*′*^*V* ^*β′*^, where *α*^*′*^ = 0.0026, *β*^*′*^ = 1.18 ± 0.16. This scaling relation, while slightly shallower than the above values of *β*, is consistent with the intraspecific metabolism exponent for *M. leidyi* obtained by Kremer [72], which would indicate that the metabolic cost of spawning scales approximately linearly with volume.

## Discussion

We have described a multigenerational lineage, namely the MV31 strain of the lobate ctenophore *Mnemiopsis leidyi* housed in an inland laboratory culture system. Few reports exist of sustained ctenophore cultures, and this is the first measurement of their fecundity after multiple generations raised in captivity. We show that a mixed zooplankton culture of the copepod *Apocyclops panamensis* with the rotifer *Brachionus rotundiformus* provides a convenient and nutritious diet for the ctenophore *M. leidyi*. The protocol presented here allows year-round culture independent of culturing facilities for fish, shrimp or wild-caught zooplankton.

The observed sizes of this strain of the inbred lab-grown ctenophore, as well as its spawn size, tends to be somewhat less than that of some but not all wild-caught animals. The viability of spawn, as defined by the ratio of hatched cydippid larvae to total spawned embryos is, on the other hand, comparable to that of wild-caught animals. While inbreeding can reduce fitness of a population to conditions outside of the labs, the MV31 strain has likely undergone sufficient selection over multiple generations to favor viability in lab conditions.

We have not recorded life spans of *M. leidyi* adults. We replace populations once they physically start to deteriorate or if there is an extended period of poor egg counts, which tends to happen after about 4-6 months. Our pseudokriesel systems likely produce some degree of turbulence and shear, and adults occasionally contact the bottom surface and the mesh screens, as a result of which they may fail to realize their life expectancy in the wild.

The protocol described here may be further optimized to use minimal human and material resources by systematically studying the impact of varying various culturing parameters. For example, the extent of improvement in egg production and hatch rates offered by including *Apocyclops panamensis* in the ctenophore diet could be compared to a baseline diet of only rotifers. Some culturing parameters to consider optimizing include rotifer feed and harvest rates, intensity and wavelength of light cues for ctenophores, the composition of the feed provided to the rotifer-copepod co-culture, and the population densities of adult ctenophores in pseudo-kreisels as well as those of larvae in static grow-out containers. Facilities that require maintenance of multiple ctenophore lineages, for example due to genome modifications, will need to explore system designs that obviate the risk of cross-fertilization across strains due to broadcast spawns that release sperm and unfertilized oocytes. The inland laboratory culture system as described is capable of sustaining longterm multigenerational cultures, satisfying an essential step needed to maintain a model ctenophore system.

## Methods

### Culturing Protocols

All cultures are maintained in artificial sea water (ASW) at a salinity of 26 parts per thousand (ppt), made by mixing Instant Ocean® sea salt in deionized water. Reproductive adults are kept in two 340 L pseudokriesels, with about 25–50 animals per tank, and fed with a mixed zooplankton culture at regular intervals through an automated peristaltic pump. A detailed description of the aquarium components is provided in Supplementary Materials Section S1 and Fig. S4.

Mixed multigenerational cultures of *Brachionus rotundiformis* (originally sourced from Reed Mariculture) and the copepod *Apocyclops panamensis* (Apex-Pods™, originally sourced from Reed Mariculture) are grown in 20 L buckets, and maintained at a temperature of 30(2) °C. The mixed culture is fed with mixed algae concentrate “RG-Complete” (Reed Mariculture). With culture volumes of 12–16 L, an algae feed rate of 12–14 mL/d, and with a harvest rate of 3.0–3.6 L/d, we have observed population densities of 200–500 ind/mL for rotifers, 1.0–2.5 ind/mL for copepod nauplii, and 1.0–2.5 ind/mL for copepods and copepodites in these mixed cultures. Compared to the protocol of Presnell et al. [6], we use a more intense rotifer culture with greater density and RG-feed rates. Populations are rebalanced occasionally by manual harvesting to reduce the density of copepods.

We also maintain two additional supporting cultures, one for the haptophyte *Isochrysis galbana*, as well as pure cultures of *Apocyclops panamensis* (see foodchain in Fig. S1). However, we have not observed difficulties with growing ctenophores in the absence of these supporting cultures. Detailed culturing procedures are described in Supplementary Materials Section S1 and Figs. S2 and S3.

We report the populations of rotifers and copepods observed in each of our four mixed zooplankton cultures over several days in Supplementary table S2. A 1 mL sample of each culture was transferred to a Sedgewick-Rafter counting chamber and a drop of vinegar was added to immobilize rotifers and copepods. Populations of rotifers were then recorded. A 10 mL sample was also taken in a glass dish, to which a few drops of vinegar were added. Upon cessation of swimming, populations of copepod nauplii were recorded. Separately, the populations of copepods and copepodites were also recorded from this sample.

### Collecting Spawns and Rearing Larvae

Ctenophores are kept in a light-controlled room and a full-spectrum LED light (MARS aqua, MZAQ-300-100LED) is used to provide light-dark cues for spawning, with a dark period of 16–18 h. Similar to the wild Floridian population of *M. leidyi* from which this lineage is derived, the animals spawn 4 hours post darkness (hpd), first releasing sperm and then eggs [6, 73]. To collect the spawn, animals are scooped out of tanks just prior to 4 hpd into 150 mm-wide Pyrex dishes, and left undisturbed for 1–2 h. Small 75 mm dishes are then used to scoop out adults which are then returned to their tanks.

Healthy larvae hatch between 18–24 hours post fertilization (hpf). To sustain them, we filter our mixed zooplankton culture through a 100 µm sieve (Fisherbrand 352360) to remove copepods and copepodites, adding enough of it to produce a rotifer density of 1–10 ind/mL in the bowl. After 3–5 days, larvae begin to be visible to the unaided eye and are subjected to a water change. A fresh bowl of ASW is seeded with filtered zooplankton at the above density, and larvae are transferred using a disposable plastic transfer pipette (e.g. Fisherbrand 137119AM). Subsequent transfers of juveniles are done every 5–7 days and fed with unfiltered zooplankton culture. Depending on the feed rate, *M. leidyi* larvae mature into adult lobates within 3 weeks. Maturation rate can be shortened to ∼ 14–16 days with enhanced feeding rates or prolonged with reduced feeding rates.

### Quantifying Reproductive Output

Culturing data was collected in June-July 2023. During this period, we maintained two independent pseudokreisels, tank “A” containing 45 animals and tank “B” containing 41 animals. All individuals came from spawns in late April 2023, and were introduced to tanks A and B on May 25th and June 13th respectively. Over a period of 10 days between June 13, 2023 and July 15, 2023, 30 individuals were sampled from tank A, with three animals studied each day. Further, over 4 days from July 17th-20th 2023, 16 individuals were sampled from tank B, with four animals studied each day. The two tanks comprised all available animals being maintained under the described protocol. Since individual lobates are difficult to distinguish from one another, ctenophores were sampled with replacement each day.

Lobate adults were selected randomly and placed individually in 150 mm bowls for 1.5 h from 3.5–5 hpd and allowed to spawn. Care was taken to fill all bowls in the study with similar volumes of water, approximately 0.5–0.6 L. For each animal, we used a compass and ruler to make measurements of the oral-aboral length, lobe-tip to aboral length, and the maximum width at the mouth in the pharyngeal (stomodaeal) plane. Following the spawn, the adults were removed carefully with a small dish, minimizing the amount of water removed and avoiding disturbing any settled eggs. The number of eggs in the dish, as well as hatched larvae the following day, were then counted as described below.

To enable counting of eggs and larvae, markings were traced onto the spawn dish to divide it into 30 equal-area segments using the grid shown in Fig. S5. Counts were made under a dissection microscope at 6x magnification, scanning depth-wise in each segment, and then moving to the neighboring segment in the pattern shown in Fig. S5. Rotation of the bowls was avoided, which would otherwise move embryos and larvae over region boundaries. Counts were recorded for all individual segments, and the totals are reported in Tables S3 and S4. All eggs, fertilized and unfertilized, were counted on the day of the spawn. The next day, at approximately 27 hpf, the number of successfully hatched larvae was counted. A small number of larvae are sometimes observed that are unusually small and remain close to the bottom of the dish. These larvae generally had developmental defects which precluded their survival to adulthood, and were therefore excluded from the count of successfully hatched larvae.

A summary statistic of successful hatch rate was calculated by plotting number of larvae against the number of eggs and fitting a linear model with zero intercept to the values compiled from both tanks. The best fit model was constructed by minimizing the mean absolute error, rather than mean squared error, in order to reduce the effect of outliers. The 95 % confidence interval of the slope was estimated by bootstrapping with 10,000 random samples of the same size as the dataset, chosen with replacement.

### Modeling the size and reproductive output of *Mnemiopsis*

To analyze scaling laws in healthy and fertile individuals, in all datasets, unsuccessful spawns defined as having <25 eggs were removed from analysis. When literature data was not tabulated but only plotted, WebPlotDigitizer was used to extract data points [74]. To enable comparisons across separate studies and to make a connection to metabolic models, we converted all size measurements to volume, through reasonable modeling assumptions described in the Results.

The animals analyzed by Reeve et al. [64] were grown for a period of 16–18 days at food concentrations of either 100 or 200 copepods/L. Three time series of data were analyzed in this paper (200H, 100I, and 200I in [64]), where the number of spawned eggs was measured *vs*. time, as well as the dry mass in µg carbon for the corresponding ctenophore, again *vs*. time (i.e. at several different times).

The length measurements made in the present study included the lobe-tip to aboral length *L* and the lobe-lobe width *D*. We calculated the volume as *V*∝ *D*^2^*L*, with a proportionality constant of *π/*4, corresponding to a cylindrical approximation. It is worth noting that the numerical prefactor does not affect the scaling relations analyzed in the text. To compare with literature data when only length was measured, we examined the scaling of *D* with *L* for our sample to obtain a scaling relationship of the form *D* ∝ *L*^*d*^. We then used the resulting best-fit equation to infer D from L, and hence calculate the volume from *L* alone. To obtain the predicted volume from the dry mass, the following method was used. In experiments A and B of [64], the authors provide length data along with dry mass in µg carbon. Fitting this data with a scaling relationship of the form *L* ∝ *m*^*l*^ between length *L* and dry carbon mass *m*, and together with the relationship between length and volume described above, we obtain the relationship between volume and dry mass. Thus, for a given dry mass, the length and therefore the predicted volume could be obtained. One minor complication was that for the plotted data in [64], the times at which egg production was measured did not coincide with the time points at which carbon mass was measured. In these cases, a linear interpolation was used between the two neighboring time points to obtain an estimated value of dry mass at the time of spawning.

The parameters for all scaling relationships were estimated by linear regression between the base-10 logarithms of the two quantities. Uncertainty in the slope was estimated under the assumption of normality of the residuals. Numerical analyses and plots were done using the NumPy, SciPy and Matplotlib packages for Python.

## Acknowledgements

The authors would like to thank Eric Henry of Reed Mariculture, Mackenzie Neale and the aquarists at the Vancouver Aquarium, Josh Wagner at the Long Beach Aquarium of the Pacific, and Jim Stime Jr. of Jelliquarium, all for advice on aquaculture and system design. P.G. has received support from a MITACS fellowship. There are no PG numbers to report for this project. We would like to thank the numerous graduates and undergraduates who were captivated by this project and devoted their time by volunteering to set up cultures, trouble-shoot protocols, build and assemble equipment, and by contributing to culture upkeep and maintenance. These people include Lana Kashino, Kyra Boulding, Jacqueline Lin-Zheng, Anna Zhitnitsky, Gabriel Dall’Alba, and Charlotte Barclay.

## Author contributions

P.G and S.S.P developed the culturing protocol in consultation with W.E.B, building on previously established methods from W.E.B’s lab. C.F counted and quantified spawn outputs and densities of feed cultures. P.G and S.S.P. wrote the manuscript with editorial input from all authors. S.S.P conceived and supervised the project.

