## Supplementary Materials for "Reproductive success of inbred strain MV31 of the ctenophore *Mnemiopsis leidyi* in a self-sustaining inland laboratory culture system"

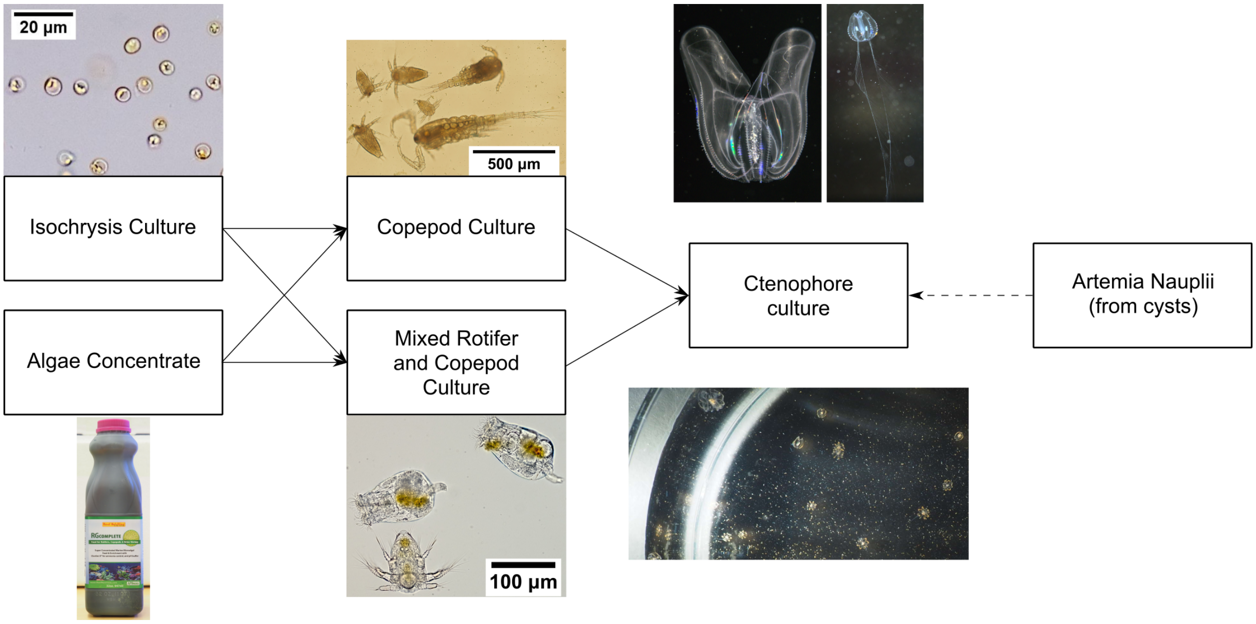

Figure S1: Overview of the foodchain. An algae concentrate, RG-Complete (Reed Mariculture), supplemented with live *Isochrysis galbana* is fed to mixed rotifer and copepod cultures, as well as (optionally) copepod-only cultures. The image of the copepod culture sample shows various life stages of *Apocyclops panamensis*. Two rotifers and a copepod nauplius are pictured in a sample from the mixed rotifer and copepod culture. The ctenophore culture images show a lobate adult (upper left, approx. lobe-to-lobe width 2.5 cm), a young cydippid larva with extended tentacles (upper right, approx. image width 10 mm), and several larvae in a 150 mm-wide bowl, some with recently formed lobes (bottom, approx. larvae size 2–10 mm). *Artemia* nauplii are not a core component of the food chain and are only used in the event of a crash of the mixed rotifer and copepod culture.

### S1 Detailed Culturing Protocols

*Mnemiopsis leidyi* (hereafter abbreviated to *Mle*) are raised on a live zooplankton diet, and cultures of these prey animal are in turn fed with a mix of live algae and algal concentrate (Fig. S1). All animals are maintained in artificial sea water (ASW). The equipment and procedures for the various cultures are described in detail below.

#### S1.1 Preparation of Artificial Sea Water

Deionized water is prepared using a filtration system consisting of a 1 µm sediment filter, two carbon block filters, reverse osmosis membranes, and a deionization chamber containing mixed anion and cation exchange resins. Tap water available in our lab has 13 parts per million (ppm) of dissolved solids, which alternatively allows for direct use, bypassing the reverse osmosis and deionization filtration stages, provided Prime® (Seachem) is used to condition the water. The filtered and/or treated water is collected in a 200 L barrel, and Instant Ocean® sea salt is added slowly while mixing the water using a submersible pump. Salinity is adjusted to 26 parts per thousand (ppt), measured using a refractometer. The water is allowed to mix for at least two days before being used.

#### S1.2 Culturing protocol for *Isochrysis galbana*

While none of our zooplankton cultures described later depend crucially on live algae, we maintain a culture of *Isochrysis galbana* (Parke 1938) (Parke, 1949) to supplement the algae concentrate as a food source for zooplankton cultures. Our *Isochrysis* strain was sourced from the Canadian Center for the Culture of

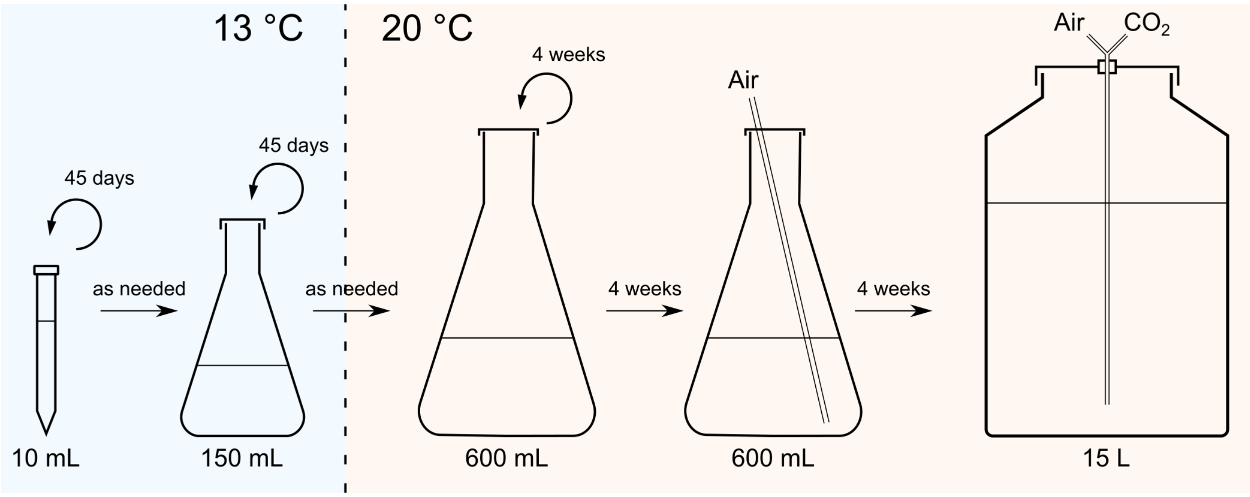

Figure S2: Schematic for phytoplankton culture. The quantities listed at the bottom are culture volumes, not container volumes.

Microorganisms (isolate 633). *Tisochrysis lutea*, also known as the T-Iso (isolate 601) may in some cases be preferable to *Isochrysis galbana* due to its nutrient profile and tolerance to a higher temperatures (Mai *et al.*, 2021). We have found both algae cultures suitable, but have not made a systematic comparison between them.

##### S1.2.1 Equipment and its Arrangement

All algae cultures are grown in Guillard's F/2 medium (Guillard, 1975). This is prepared by adding 135 µL of each of parts A and B of a growth medium concentrate, namely "F/2 Algae Food" (Fritz Aquatics), for every 1 L of ASW. The ASW used for this purpose can be adequately sterilized simply by microwaving (Keller *et al.*, 1988); we microwave 6 × 600 mL of ASW in six Schott bottles for 12 min at 900 W. The ASW is allowed to cool before adding the nutrient concentrate solutions. Air tubes supplying algae cultures are fitted with 0.2 µm PTFE filters (Fisherbrand 13100115).

The production culture is maintained at room temperature (≈ 20 °C) in a 6 gal polyethylene terephthalate (PET) carboy container (FerMonster™), with a culture volume of approximately 15–18 L. The culture is bubbled with air. Optionally, carbon dioxide can be mixed into the air supply, with its rate controlled by a timed solenoid valve as well as a pressure regulator, and calibrated to produce the optimal pH (Kaplan *et al.*, 1986).

The production culture is supported by a series of auxiliary cultures of various volumes (Fig. S2). A 10 mL culture is maintained in a 15 mL conical centrifuge tube (Corning™ 430791), and 150 mL cultures are maintained in 250 mL conical flasks; both are placed in a 13 °C incubator. A room temperature culture of 600 mL is maintained in a 1 L conical flask and used to seed each new production culture. This culture is bubbled about 5 days prior to being used as a starter culture to increase its population.

Room-temperature cultures are continuously illuminated by a full-spectrum visible light LED light source (MARS aqua, MZAQ-300-100LED) at minimum working intensity placed about 20 cm away from the cultures, and producing about 10,000–20,000 lx of illuminance. It may be preferable to grow the production culture with a light-dark cycle such as 18 h light and 6 h dark, rather than 24 h of light, however we have not optimized this aspect. The 13 °C cultures are weakly illuminated at ≈100 lx by a red and blue LED lamp. The lower temperature and weak illumination cause these latter cultures to grow slower and require less frequent upkeep while still serving as backups against crashes of the room-temperature cultures.

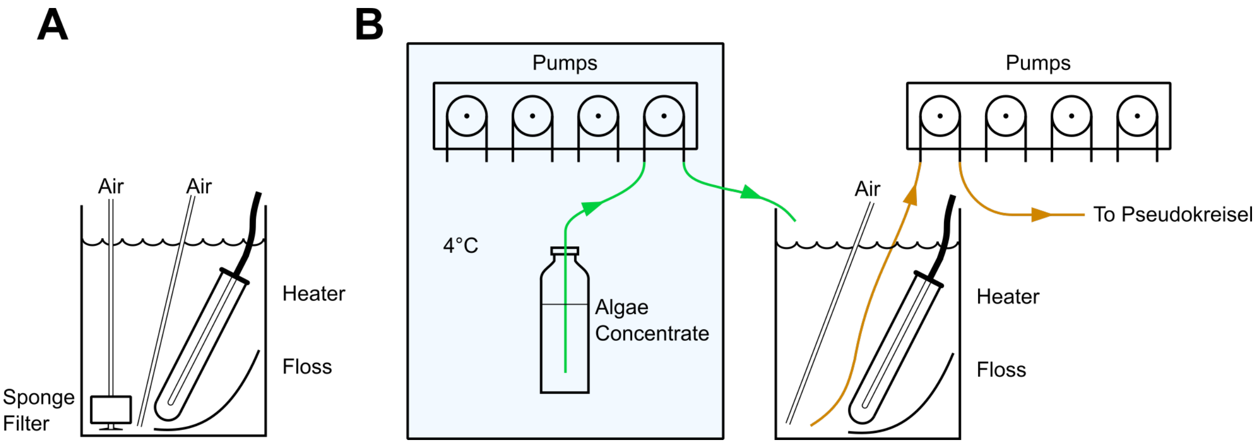

Figure S3: Schematic for zooplankton culture. Copepod cultures (A) and mixed cultures of rotifers and copepods (B) are grown in 20 L buckets. The latter are automatically fed and harvested using peristaltic pumps (Jebao DP-4). One culture of each type is shown.

##### S1.2.2 Starting and Propagating Algae Cultures

Cultures are always started in sterile containers. The 6 gal carboy and its submerged air line are sterilized in bleach water (1:500 dilution of household 5 % bleach). Bleach water is drained out and the carboy is rinsed with dilute sodium thiosulfate in ASW (≈5 mL of 0.2 M $Na_2S_2O_3 \cdot 2\,H_2O$ in 600 mL of ASW) before being filled with freshly microwaved warm ASW (<60 °C) and bubbled through the air line. Concentrated nutrients and starter culture are added the following day. The culture takes 10 days to reach its maximum sustainable population and can then be harvested. We harvest 3.6 L of the culture each day and replace the harvested volume with fresh media. The culture is continued for approximately 4 weeks after which it generally needs to be discarded due to the inevitable and visible growth of contaminating organisms on the inner surface of the carboy, accompanied by a reduction in intensity of the yellow-green colour of *Isochrysis*.

The 600 mL auxiliary culture is propagated at the rate set by the production culture. Approximately every 4 weeks, a small amount of the current auxiliary culture is used to seed a fresh culture of the same volume, and subsequently bubbled for 3–6 days to increase its density before being used to seed the next production culture. The production culture can be seeded either directly at the target volume of 15 L or at a lower volume of ≈10 L, and then gradually increased in volume once the algae density increases, by adding 3.6 L of sterilized ASW each day.

Backup cultures at 13 °C are propagated every 45 days. The 10 mL and 150 mL cultures are normally maintained independently, only starting new cultures of their respective types, but they can each be used to start a new culture of a larger volume to aid recovery from crashes. All cultures that are maintained without bubbling are only opened and propagated while working close to a Bunsen burner to reduce chances of contamination from air-borne organisms.

Mature cultures present a deep yellow-green to brown colour which varies seasonally with ambient conditions. We have not standardized culture densities.

#### S1.3 Protocol for Culturing *Brachionus rotundiformis* and *Apocyclops panamensis*

The essential food for our *Mle* cultures is a mixture of the rotifer *Brachionus rotundiformis* with the copepod *Apocyclops panamensis* (both sourced from Reed Mariculture). In addition to the mixed cultures, we maintain pure cultures of *Apocyclops panamensis*, mainly as backup to replenish copepod populations in the mixed cultures in the unlikely event of a collapse. We often find however that the population densities of copepods in the co-cultures exceed those in the isolated populations of *Apocyclops panamensis*.

##### S1.3.1 Equipment and its Arrangement

Our culture equipment for the mixed cultures consists of 20 L cylindrical buckets containing a 75 W immersion heater and a rigid air-line (Fig. S3B). The air tube is tipped with a small piece of flexible tubing, which is easier to clean when it gets clogged with salt precipitates and detritus. The air tube is also weighed down using a rubber stopper to remain submerged. A piece of rotifer floss (Reed Mariculture), approximately 20 cm × 20 cm is placed in the bucket, typically placed underneath the heater. This arrangement is a simplification of the "Compact Culture System" recommended by Reed Mariculture and reduces effort during daily maintenance.

Co-cultures are maintained in 26 ppt ASW at 26–30 °C, with an airflow sufficient to produce churning throughout the bucket (the air pump is a Marina 200, rated $110\,\mathrm{L\,h^{-1}}$). The cultures are primarily fed with RG-Complete (Reed Mariculture) which is a concentrated mixture of algae. Supplementing with live *Isochrysis galbana* algae is optional, but we believe it improves rotifer density and culture stability. Under usual operation, our cultures have a volume of 12–18 L and are fed with 10–14 mL of RG-Complete and 0–0.6 L of live algae culture per day. A total of 2.4–3.6 L of culture is harvested each day from each of 4 cultures to feed ctenophores. With this arrangement, our cultures contain $150\text{–}400\,\mathrm{mL^{-1}}$ of rotifers, $1\text{–}3\,\mathrm{mL^{-1}}$ of copepods or copepodites, and $2\text{–}10\,\mathrm{mL^{-1}}$ of copepod nauplii. Table S2 shows measured population densities in our four cultures over a period of about 2 weeks. Automated dosing pumps (Jebao DP-4) are used both for harvesting rotifers and for adding RG-Complete at regular intervals spaced evenly throughout the day. The addition of ASW to compensate harvested volume may also be automated, which minimizes the maintenance necessary to maintain roughly constant water volume, although this results in additional wear and tear on the pumps. For the data collected in this study, manual daily addition of ASW was implemented.

Pure cultures of *Apocyclops panamensis* are placed away from the mixed cultures to reduce opportunities for contamination. These cultures are maintained in 20 L buckets, with a sponge filter which also aerates the culture, and a 75 W heater, at the same salinity and temperature ranges as the mixed cultures (Fig. S3A). In steady state, our pure copepod cultures have a volume of 9–12 L and are fed manually once a day with 1–5 mL of RG-Complete and optionally 0.5–1 L of live algae culture. Care is taken to avoid overfeeding—we ensure that the greenish colour from the previous feeding has dissipated before more food is added. Anywhere from 0–1 L of culture is harvested each day, and the volume is replaced with some live algae culture if available, or ASW. In this state, about 1–4 copepods or copepodites and 0–3 nauplii are present in 1 mL of culture. In our experience copepod populations vary considerably over time, and are slow to recover following water changes or crashes.

##### S1.3.2 Maintenance

To maintain a reliable supply of live zooplankton, their culture needs daily maintenance and monitoring. Detritus from the food sources collects on all surfaces and needs to be removed from cultures each day. To do this, the rotifer floss present in each culture, which has already accumulated detritus, is used to wipe down all submerged surfaces. Any large pieces of floating detritus may be manually filtered by passing the culture water through the rotifer floss. The rotifer floss is then thoroughly rinsed under a jet of tap water in the sink, before being wrung dry and returned to its culture bucket. The density of rotifers is monitored visually, simply by turning off the air flow and shining a bright light through the bucket walls. A more rigorous count is occasionally recorded using a Sedgewick-Rafter counting slide. Optionally, the cultures can be subjected to a complete water change about once every two months, however, with experience, water changes can be avoided unless culture conditions deteriorate (see troubleshooting in Section S1.3.3). To perform a water change, tubes and heaters are disconnected, and the culture is passed slowly through a 40 µm filter, preceded by a piece of rotifer floss to catch and filter larger detritus clumps. The bucket, heater and tubes are wiped and rinsed, and then the collected rotifers are added back along with the appropriate volume of ASW.

Over time, the relative populations of the two species have been observed to shift in favour of the copepods. When a population imbalance is observed during routine visual inspection (typically about once a month), the population of reproductive copepods is reduced by passing a fraction of the culture slowly through a 120 µm filter. The copepods caught in the filter can be manually added to the ctenophore pseudokreisels. This task may be strategically performed the day prior to embryological experiments to boost spawn production in the ctenophores.

Pure cultures of *Apocyclops panamensis* are similarly monitored visually each day. We replace the water in these culture about once every two months, along with cleaning the sponge filter to reduce built up detritus.

Samples of the culture are occasionally inspected under a microscope for the presence of contaminant rotifers (see troubleshooting in Section S1.3.3).

##### S1.3.3 Troubleshooting and Restarting Rotifer/Copepod Co-cultures

During daily inspection, several properties of the co-culture are observed for deviations from the norm. The colour of the culture should be pale yellow and rather transparent, with a light green tint following a recent addition of algae concentrate. A deeper green indicates that more food is being added than consumed. The odour, which is usually earthy, may turn foul if overfeeding persists due to decay of excess algae concentrate. Density of rotifers and copepods can also be easily observed directly in the bucket once air flow is temporarily stopped and the bucket is illuminated through the walls – adult copepods are larger and make quick sporadic movements, compared to rotifers which are smaller and move on smoother trajectories.

While routine inspection with the unaided eye is usually sufficient to notice and correct deviations from the steady production of zooplankton, a closer inspection may be warranted if difficulties persist. A sample of the culture is taken into a Sedgewick-Rafter counting slide and observed through a dissection microscope at (6×)–(20×) magnification. After inspection of live zooplankton, a drop of vinegar is used to kill the animals to ease counting. Potential early indicators of poor culturing conditions include slower or inhibited swimming in rotifers, and reduced ratio of female rotifers carrying eggs on their posterior over the total number of females (Hoff *et al.*, 2007). The presence of decaying matter in the culture may also be inferred from an increased number of ciliates. To correct these problems, first one should ensure that there is no excess feeding, and any excessive food and decay products should be removed by doing a water change. Starting from a clear filtered culture, the feeding rate should be gauged based on how quickly the colour from a feeding is cleared, and gradually increased over the next few days. The volume of the culture should be increased in proportion to rotifer counts, maintaining about 100–300 $mL^{-1}$ during the recovery period. Harvesting can be started and gradually increased over 2–3 days once the culture reaches the target volume. If harvesting cannot be initiated within about 7 days of restarting a culture, a water change will be necessary.

It is best to maintain multiple rotifer cultures so that a collapse of one can be corrected simply by seeding a new culture from a sample of another healthy culture, rather than relying on a smaller population of healthy reproductive adults that may be present in the failing culture alongside a potentially bigger number of dying rotifers. In practice we keep 3-4 culture buckets active at any given time.

In the event that a mixed culture has too few copepods, this can be corrected by adding a portion of a pure *Apocyclops panamensis* culture to it. It can take as long as a month for the copepod population to recover due to their slower life cycle.

If a pure *Apocyclops* culture is found to be contaminated with rotifers, the culture may be filtered through a 120 µm screen. This process retains adults and larger copepodites, but allows nauplii and rotifers to pass through, which are then either discarded or concentrated and fed to the ctenophores. A new culture is started from the adults. This culture takes about 4 weeks to reach steady state again due to the slower life cycle of this species and must not be harvested during recovery. To recover from a complete collapse of pure copepod-only cultures, large copepods in the rotifer cultures can be harvested using a 120 µm filter. It is likely that such a culture will have some contaminant rotifers and may need further corrective measures if a pure copepod culture is desired.

#### S1.4 Culturing Protocol for *Mnemiopsis leidyi*

##### S1.4.1 Equipment and its Arrangement

Populations of lobate *Mle* individuals are maintained in 90 gal pseudokreisels (Jelliquarium) (Fig. S4). While all fresh ASW is prepared at 26 ppt, due to evaporation the salinity stabilizes at 29–30 ppt, for our rates of feeding and water replacement. Circulation of water in the tank is maintained by a 1/12 HP pump (Iwaki MD-40RT-115) at around 1.0 RPM, which ensures that all animals are kept in motion, minimizing “dead zones” where animals can stay stationary. An in-line chiller (Tradewind IL-15-S) maintains a temperature of (19 ± 1) °C. The main section of the tank is separated from the refugium area containing the exit tube by a 400 µm screen. A finer mesh can more easily become clogged with detritus, disrupting the flow. The tank is connected to a sump area containing equipment to maintain water quality. A wet-dry filter containing “bio-balls” and a submerged “bio-brick” provide surfaces for nitrogen-processing bacteria to grow in aerobic

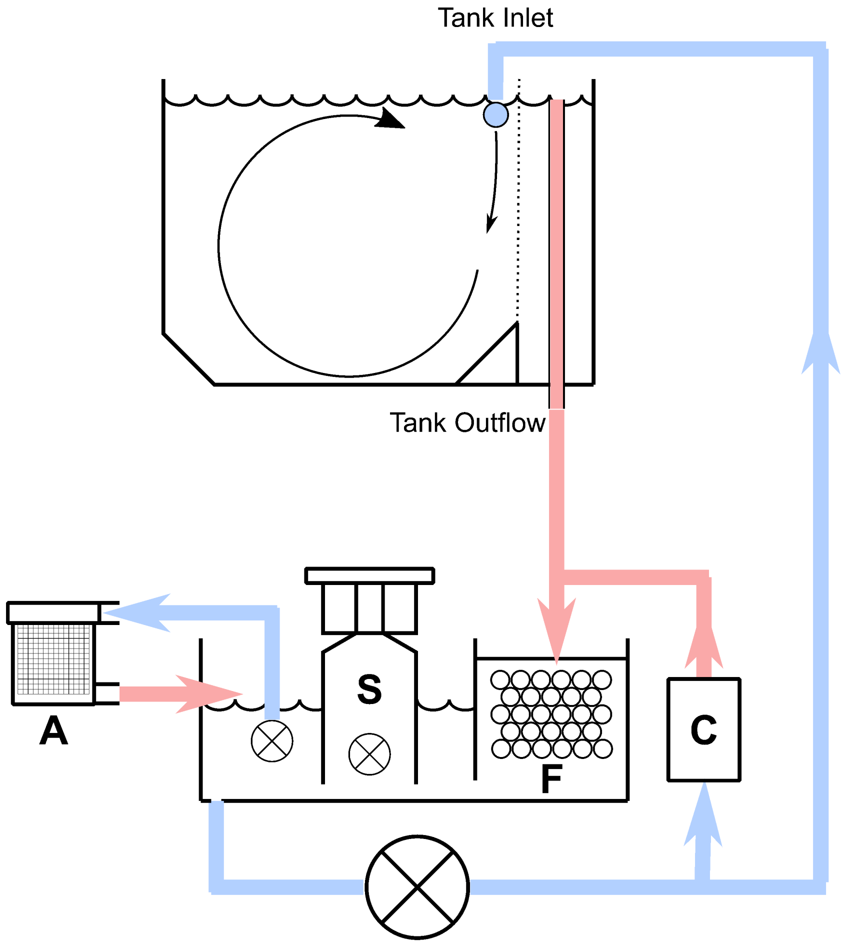

Figure S4: Schematic for ctenophore pseudokreisel systems. The pseudokreisel (above) contains ctenophores and is connected to a sump (below). The dotted line in the pseudokreisel indicates screens separating the refugium which contains the drain. The tank inlet is connected to a spray bar that maintains a laminar flow downwards over the screens. Water moves away from the pseudokreisel and towards the sump along the red paths and moves away from the sump and returns to the pseudokreisel along the blue paths. A: algae scrubber, S: protein skimmer, F: wet-dry filter, C: in-line chiller, ⊗: pumps.

conditions. A protein skimmer extracts organic compounds and detritus. An algae scrubber provides a surface for macroalgae to grow, extracting nitrates and phosphates from the water in the process and reducing growth of photosynthetic organisms on surfaces in the main section of the tank.

The pseudokreisels are kept in a light-insulated room, and illuminated by a full-spectrum visible light LED light source (MARS aqua, MZAQ-300-100LED, normally used for coral growth). The lights are powered on a timer, with a 16 h dark period and an 8 h light period which ends at 6 AM, allowing collection of spawn at 10 AM, four hours after lights-out. The timing of the spawn can be adjusted to one's preference; we find in practice that the animals adjust to new times within a couple days. The 4 h interval from darkness to spawn is characteristic of the Floridian population of *Mle*, which this lab lineage derives from. The "Northern population" found near Woods Hole Massachusetts has been found to spawn 8 hours post darkness (Pang and Martindale, 2008).

Cydippid stage *Mle* individuals are raised in either 150 mm-wide bowls or small static aquariums containing 2–8 L of ASW (Section S1.4.3).

##### S1.4.2 Maintenance

Pseudokreisel tanks are automatically fed with the mixed culture of rotifers and copepods at regularly spaced intervals throughout the day, using peristaltic pumps on a set timing schedule. The resulting rise in volume of water, as well as the separate issue of the build up of detritus, are both managed by manually siphoning out water from the bottom of the main section of the tank daily. Waste water is kept in buckets for one day after $\approx 1$ mL/L of bleach is added, to comply with biosafety guidelines for this invasive species. Protein skimmers are emptied each day, and the skimmer waste water collection rate is adjusted to ensure that a consistent volume of around 100–200 mL/day is collected, varying slightly with feeding rates and tank cleanliness. Macroalgae in algae scrubbers is removed every two to three weeks by manually scraping algae growth off of the algae scrubber screens.

The various surfaces in the main section of the tank are cleaned about every other month. The screens separating the main pseudokreisel from the refugium are cleaned monthly using rotifer floss or a similar scrubber. The other rigid surfaces (walls and floor of the tank) are wiped with a rubber squeegee, the suspended detritus is allowed to settle, and subsequently detritus is siphoned out before restarting the flow and reintroducing the animals. Frequent observations of an imbalance in water levels between the main tank and refugium indicates that a more thorough cleaning of the screens is required, which is implemented as follows. With all animals removed, and circulation is stopped, and the screens are subjected to a strong air pressure through tubing connected to a high pressure air supply. This air bubble-cleaning procedure effectively dislodges detritus and photosynthetic organisms from the mesh.

The concentration of ammonia and nitrites in aquariums must be kept below $\approx$0.2 ppm, and is confirmed by water tests. In mature (homeostatically equilibrated) tanks, frequent testing is not necessary. Elevated levels of nitrates and phosphates can increase the growth of photosynthetic organisms on illuminated tank surfaces. The screens separating the refugium in particular are sensitive to this − growth on this surface blocks the mesh holes and leads to disruption and heterogeneity of the aquarium flow. In this situation, ctenophores can become trapped against the screens in local regions of high flow, which rapidly leads to tissue damage and disintegration.

Feeding amount and frequency must be balanced with the population and size of the ctenophores in the tanks. Overfeeding is a likely cause for elevated nitrogenous waste products in the aquarium. If fouling of water is observed or suspected, a high proportion (as much as 80 %) of the aquarium water can be replaced with fresh ASW. Adults can survive several days with little to no feeding. While water quality recovers, a manual directed feeding approach using pipettes can be employed, where concentrated zooplankton are released close to the lobes and mouths of adults. We have rarely found this to be necessary, however.

In our observations, healthy adults have open lobes, directed slightly outward from the mouth along the pharyngeal axial plane (see Fig. S1 upper right). When food is added to a food-depleted tank, the guts of healthy adults become visibly occupied in minutes. The integrity and length of the lobes and auricles are also key indicators of health (as noted by Presnell *et al.* (2022)). In our experience, a population of adults can easily live for at least 6 months, after which the numbers start to decrease and a new generation of larvae should be raised for replacement.

###### S1.4.3 Collecting Spawns and Rearing Larvae

At approximately 3.5 hours post darkness (hpd), healthy and gravid lobate Mle adults are removed from their pseudokreisel habitat and placed into 150 mm-wide crystallizing dishes (PYREX 3140150). They are left undisturbed for 2–3 hours to spawn. Multiple adults can be spawned in one bowl – we have spawned as many as 15 at a time in one dish, although overcrowding is likely to be detrimental (Ramon-Mateu *et al.*, 2022). Sperm are released first, around 4 hpd, followed by eggs which are released slowly over the next hour. The adults are then removed from the dishes using a small bowl, leaving behind most embryos in the dishes. Healthy adults of our lineage spawn roughly 100–700 eggs per adult in a spawn event depending on their size, averaging around 400 eggs (see Tables S3 and S4). The following day, starting around 18 hours post fertilization (hpf), larvae begin to hatch. In healthy populations, about 70 % of the embryos successfully develop into larvae (Tables S3 and S4 and Fig. 1B of the main manuscript), however, small spawn sizes $\leq 100$ embryos per ctenophore result to lower ratios of viable larvae (Fig. 1C of the main manuscript).

To sustain larvae, we filter our mixed zooplankton culture through a 100 µm sieve (Fisherbrand 352360) to remove copepods and copepodites, adding enough of it to produce a rotifer density of 1–10 $\mathrm{mL}^{-1}$ in the bowl. After 3–5 days, larvae begin to be visible to the unaided eye and are subjected to a water change, as follows. A fresh bowl of ASW is seeded with zooplankton at the same density, and larvae are transferred using a disposable plastic transfer pipette (e.g. Fisherbrand 137119AM). Water may be gently added to "grow-out" tanks by using either a drip-line or siphon consisting of a thin tube with filter paper and a pipette tip to control the flow to a slow drip. Due to the higher salinity of the water in pseudokreisels, the larvae take some time to adjust to the lower salinity in the new bowl. Subsequent transfers are done every 5–7 days and the zooplankton culture is fed as is, without filtering through the 100 µm sieve.

The cydippid stage of the animals can be extended by using a smaller amount of feed. With abundant food availability, they eventually take on the lobate form and can be transferred to pseudokreisels if desired. Lobate ctenophores have lived in static tanks for periods up to several weeks in our laboratory before being tranferred to pseudokreisels.

Due to the invasive potential of this species, untreated aquarium water should never be released into natural water bodies. We treat all aquarium water with household 5 % bleach overnight at a maximum dilution of 1:1000, before draining treated water into lab drains (leading to the sewer system). Embryos and larvae are highly sensitive to bleach, and have been observed to die within minutes at the above concentrations. Bowls used for spawning ctenophores are also rinsed into buckets where the water is treated similarly, or alternatively, such glassware is allowed to dry overnight before washing.

To enable counting of eggs and larvae under a dissection microscope, markings were traced onto the spawn dish to divide it into 30 equal-area segments using the grid shown in Fig. S5. The pattern of scanning across neighbouring segments is also shown in Fig. S5. Counts recorded in all individual segments were first totalled, and the totals were then tabulated in Tables S2 and S3 (see main text).

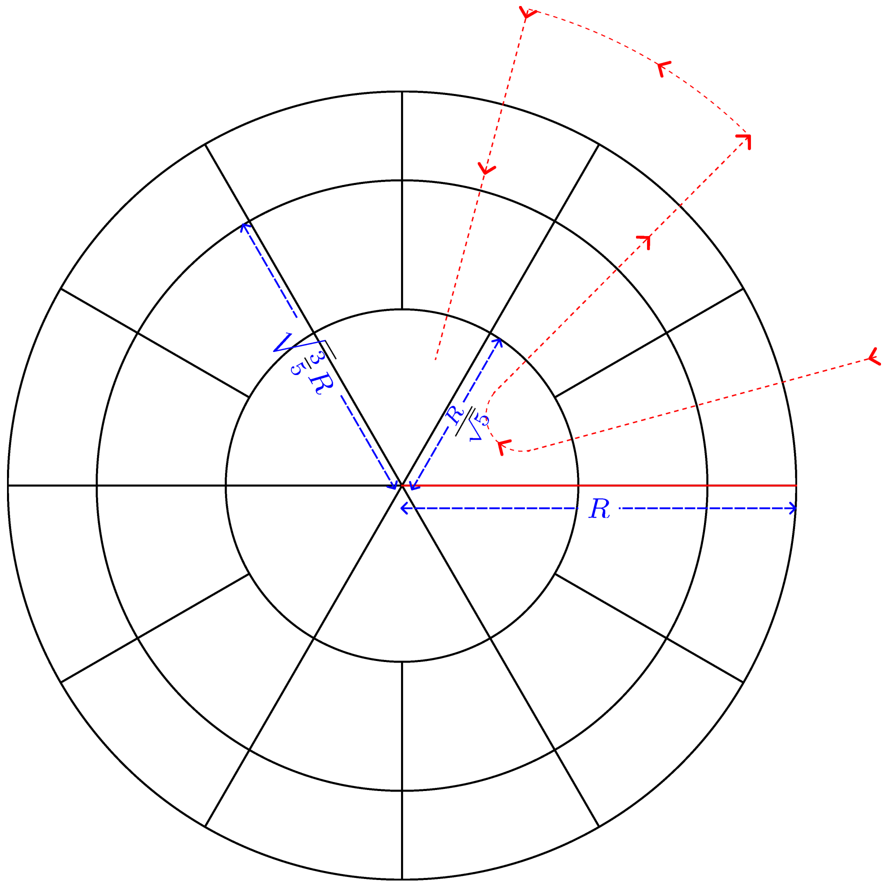

Figure S5: Grid pattern used to aid counting of eggs and larvae in bowls. This grid divides the circle into 30 equal-area regions, each of which can be seen in its entirety in the field of view of our dissection microscope. The inner diameter of the bowls was measured to be 144 mm, which sets $R = 72\,\text{mm}$. The solid red line allows identification of a starting and ending point for counting. This grid was traced onto the bottom of each glass dish using a waterproof marker. Counting was done following the path traced by the dashed red line, continuing through all regions, but without rotating the bowl.

Table S1: Measurements of reproductive output and size in the literature. All references use Floridian *M. ledyi* animals.

| **Reference** | **Growth conditions** | **Size measurement(s)** | **Tables provided** | **Number of measurements** | **Measurements with spawn size <25** |
|---|---|---|---|---|---|
| Baker and Reeve (1974) | Wild caught | Length | No | 57 | 5 |
| Baker and Reeve (1974) | First generation lab grown | Length | Yes | 51 | 2 |
| Reeve *et al.* (1989) | First generation lab grown | Carbon mass | No | 38 | 5 |
| Sasson and Ryan (2016) | Wild caught | Length | Yes | 30 | 0 |
| Present study | 31st generation lab grown | Length, width | Yes | 46 | 2 |

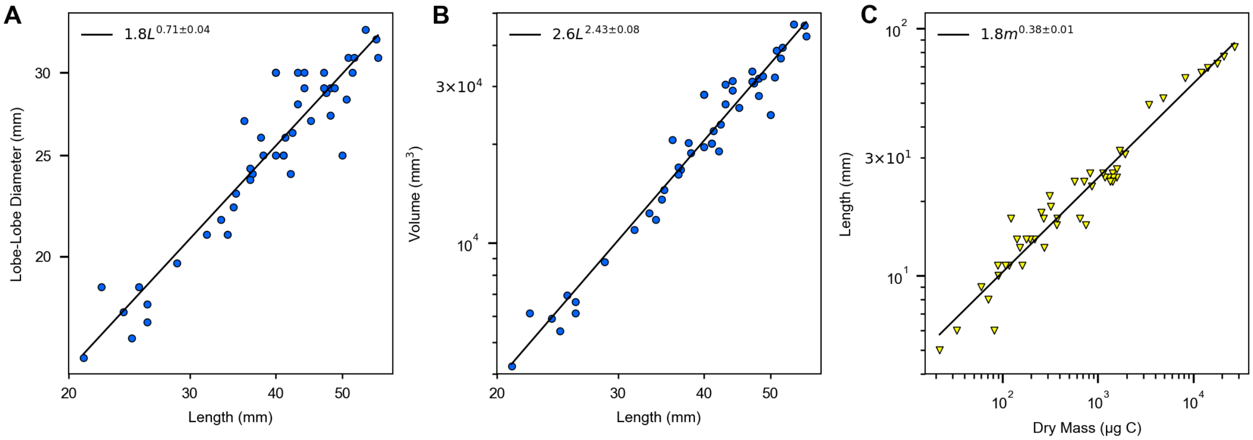

Figure S6: Scaling laws for ctenophore geometry. (A) Lobe-lobe lateral diameter $d$ of *M. leidyi* scales with aboral-lobe length $L$ as $d = (1.8\,\mathrm{mm})(L/\mathrm{mm})^{0.71\pm0.04}$. (B) *M. leidyi* volume $V$, when modelled by a cylinder of diameter $d$ and length $L$ (i.e. $V = \frac{\pi}{4}d^2L$) scales with length $L$ as $V = (2.6\,\mathrm{mm}^3)(L/\mathrm{mm})^{2.43\pm0.08}$, indicating that ctenophores become more prolate with increasing size. (C) Aboral-lobe length scales with dry carbon mass as $L = (1.8\,\mathrm{mm})(m/\mu\mathrm{g})^{0.38\pm0.01}$, supporting an approximately linear relationship between modelled volume and dry mass (see main text).

| Date | Volume (L) | Rotifers in 1 mL | Copepod Nauplii in 10 mL | Copepods & Copepodites in 10 mL | Population of Rotifers | Population of Copepod Nauplii | Population of Copepods & Copepodites |
|---|---|---|---|---|---|---|---|
| Culture S1. Fed 14 mL/day. Harvested 3.28 L/day to tank A. Temperature 30–31 °C, averaging 30.6 °C. | | | | | | | |
| 14-07-23 | 13 | 338 | 50 | 20 | $4.39\times10^6$ | $6.50\times10^4$ | $2.60\times10^4$ |
| 17-07-23 | 13.5 | 313 | 42 | 25 | $4.23\times10^6$ | $5.67\times10^4$ | $3.38\times10^4$ |
| 18-07-23 | 12.75 | 343 | 144 | 28 | $4.37\times10^6$ | $1.84\times10^5$ | $3.57\times10^4$ |
| 19-07-23 | 15.5 | 321 | 20 | 13 | $4.98\times10^6$ | $3.10\times10^4$ | $2.02\times10^4$ |
| 20-07-23 | 12 | 394 | —* | — | $4.73\times10^6$ | — | — |
| 21-07-23 | 16 | 279 | 16 | 11 | $4.46\times10^6$ | $2.56\times10^4$ | $1.76\times10^4$ |
| 24-07-23 | 15 | 303 | 11 | 9 | $4.55\times10^6$ | $1.65\times10^4$ | $1.35\times10^4$ |
| 25-07-23 | 12 | 363 | 19 | 10 | $4.36\times10^6$ | $2.28\times10^4$ | $1.20\times10^4$ |
| 26-07-23 | 13.25 | 249 | 13 | 11 | $3.30\times10^6$ | $1.72\times10^4$ | $1.46\times10^4$ |
| Culture S2. Fed 14 mL/day. Harvested 3.38 L/day to tank B. Temperature 27–28 °C, averaging 27.3 °C. | | | | | | | |
| 10-07-23 | 16 | 256 | 10 | 10 | $4.10\times10^6$ | $1.60\times10^4$ | $1.60\times10^4$ |
| 11-07-23 | 12.75 | 287 | 13 | 16 | $3.66\times10^6$ | $1.66\times10^4$ | $2.04\times10^4$ |
| 12-07-23 | 12.75 | 353 | 22 | 10 | $4.50\times10^6$ | $2.81\times10^4$ | $1.28\times10^4$ |
| 13-07-23 | 13 | 337 | 21 | 8 | $4.38\times10^6$ | $2.73\times10^4$ | $1.04\times10^4$ |
| 14-07-23 | 13 | 279 | 24 | 19 | $3.63\times10^6$ | $3.12\times10^4$ | $2.47\times10^4$ |
| 17-07-23 | 13.25 | 347 | 60 | 22 | $4.60\times10^6$ | $7.95\times10^4$ | $2.92\times10^4$ |
| 18-07-23 | 13.5 | 315 | 71 | 28 | $4.25\times10^6$ | $9.59\times10^4$ | $3.78\times10^4$ |
| 19-07-23 | 15.5 | 301 | 34 | 16 | $4.67\times10^6$ | $5.27\times10^4$ | $2.48\times10^4$ |
| 20-07-23 | 12 | 242 | 24 | 17 | $2.90\times10^6$ | $2.88\times10^4$ | $2.04\times10^4$ |
| 21-07-23 | 16 | 278 | 18 | 8 | $4.45\times10^6$ | $2.88\times10^4$ | $1.28\times10^4$ |
| 24-07-23 | 15 | 254 | 16 | 11 | $3.81\times10^6$ | $2.40\times10^4$ | $1.65\times10^4$ |
| 25-07-23 | 11.75 | 361 | 19 | 12 | $4.24\times10^6$ | $2.23\times10^4$ | $1.41\times10^4$ |
| 26-07-23 | 13.5 | 272 | 18 | 16 | $3.67\times10^6$ | $2.43\times10^4$ | $2.16\times10^4$ |
| Culture S3. Fed 12 mL/day. Harvested 3.20 L/day to tank A. Temperature 30–33 °C, averaging 31.8 °C. | | | | | | | |
| 10-07-23 | 16.5 | 297 | 37 | 9 | $4.90\times10^6$ | $6.11\times10^4$ | $1.49\times10^4$ |
| 11-07-23 | 13 | 387 | 27 | 8 | $5.03\times10^6$ | $3.51\times10^4$ | $1.04\times10^4$ |
| 12-07-23 | 12.75 | 384 | 38 | 22 | $4.90\times10^6$ | $4.85\times10^4$ | $2.81\times10^4$ |
| 13-07-23 | 13.25 | 339 | 38 | 16 | $4.49\times10^6$ | $5.04\times10^4$ | $2.12\times10^4$ |
| 14-07-23 | 13 | 404 | 35 | 14 | $5.25\times10^6$ | $4.55\times10^4$ | $1.82\times10^4$ |
| 17-07-23 | 13.75 | 389 | 39 | 17 | $5.35\times10^6$ | $5.36\times10^4$ | $2.34\times10^4$ |
| 18-07-23 | 12.5 | 383 | 45 | 14 | $4.79\times10^6$ | $5.63\times10^4$ | $1.75\times10^4$ |
| 19-07-23 | 15.5 | 330 | 44 | 13 | $5.12\times10^6$ | $6.82\times10^4$ | $2.02\times10^4$ |
| 20-07-23 | 11 | 382 | 36 | 12 | $4.20\times10^6$ | $3.96\times10^4$ | $1.32\times10^4$ |
| 21-07-23 | 15 | 287 | 27 | 8 | $4.31\times10^6$ | $4.05\times10^4$ | $1.20\times10^4$ |
| 24-07-23 | 14.75 | 328 | 33 | 23 | $4.84\times10^6$ | $4.87\times10^4$ | $3.39\times10^4$ |
| 25-07-23 | 11.75 | 369 | 35 | 14 | $4.34\times10^6$ | $4.11\times10^4$ | $1.65\times10^4$ |
| 26-07-23 | 13.5 | 286 | 35 | 8 | $3.86\times10^6$ | $4.73\times10^4$ | $1.08\times10^4$ |
| Culture S4. Fed 12 mL/day. Harvested 3.06 L/day to tank B. Temperature 28.5–30 °C, averaging 29.2 °C. | | | | | | | |
| 14-07-23 | 13.75 | 48 | 31 | 15 | $6.60\times10^5$ | $4.26\times10^4$ | $2.06\times10^4$ |
| 17-07-23 | 14 | 87 | 42 | 39 | $1.22\times10^6$ | $5.88\times10^4$ | $5.46\times10^4$ |
| 18-07-23 | 13 | 108 | 47 | 42 | $1.40\times10^6$ | $6.11\times10^4$ | $5.46\times10^4$ |
| 19-07-23 | 15 | 96 | 25 | 18 | $1.44\times10^6$ | $3.75\times10^4$ | $2.70\times10^4$ |
| 20-07-23 | 12 | 85 | 62 | 17 | $1.02\times10^6$ | $7.44\times10^4$ | $2.04\times10^4$ |
| 21-07-23 | 15 | 104 | 30 | 17 | $1.56\times10^6$ | $4.50\times10^4$ | $2.55\times10^4$ |
| 24-07-23 | 14 | 129 | 46 | 15 | $1.81\times10^6$ | $6.44\times10^4$ | $2.10\times10^4$ |
| 25-07-23 | 11 | 142 | 45 | 18 | $1.56\times10^6$ | $4.95\times10^4$ | $1.98\times10^4$ |
| 26-07-23 | 14 | 103 | 36 | 19 | $1.44\times10^6$ | $5.04\times10^4$ | $2.66\times10^4$ |

Table S2: Culturing statistics for the mixed zooplankton cultures which fed tank A and tank B. These cultures were fed RG-Complete and maintained as stated. * A dash means measurements were not made on this date.

| Date | Oral-Aboral Length (cm) | Overall Length (cm) | Pharyngeal Width (cm) | Number of Eggs | Number of Hatched Larvae |
|---|---|---|---|---|---|
| 15-06-23 | 2.5 | 5 | 2.5 | 321 | 238 |
| 15-06-23 | 3 | 4 | 3 | 296 | 245 |
| 15-06-23 | 2.5 | 4 | 3 | 407 | 357 |
| 21-06-23 | 2.1 | 3.4 | 2.1 | 217 | 156 |
| 21-06-23 | 2.7 | 4.3 | 3 | 522 | 361 |
| 21-06-23 | 2.9 | 4.4 | 2.9 | 355 | 315 |
| 27-06-23 | 3.1 | 4.7 | 3 | 682 | 294 |
| 27-06-23 | 3 | 4.3 | 2.8 | 306 | 223 |
| 27-06-23 | 2.5 | 3.6 | 2.7 | 71 | 45 |
| 29-06-23 | 2.6 | 4.2 | 2.4 | 277 | 191 |
| 29-06-23 | 1.4 | 2.1 | 1.6 | 31 | 0 |
| 29-06-23 | 2.4 | 3.8 | 2.6 | 178 | 142 |
| 04-07-23 | 2.5 | 4 | 2.5 | 430 | 361 |
| 04-07-23 | 2.9 | 4.5 | 2.7 | 531 | 420 |
| 04-07-23 | 2.4 | 3.7 | 2.4 | 657 | 511 |
| 06-07-23 | 1.7 | 2.6 | 1.8 | 40 | 0 |
| 06-07-23 | 3.1 | 4.4 | 3 | 922 | 702 |
| 06-07-23 | 3.2 | 5.1 | 3.1 | 1255 | 894 |
| 10-07-23 | 2.9 | 4.8 | 2.9 | 345 | 268 |
| 10-07-23 | 2 | 3.5 | 2.3 | 101 | 36 |
| 10-07-23 | 3.8 | 5.4 | 3.3 | 826 | 632 |
| 11-07-23 | 2.55 | 4.1 | 2.5 | 2 | 0 |
| 11-07-23 | 3.3 | 4.73 | 2.87 | 172 | 126 |
| 11-07-23 | 3.67 | 5.17 | 3 | 1234 | 834 |
| 12-07-23 | 3.13 | 4.7 | 2.9 | 518 | 384 |
| 12-07-23 | 2.9 | 4.8 | 2.73 | 483 | 291 |
| 12-07-23 | 2.55 | 4.1 | 2.5 | 416 | 204 |
| 13-07-23 | 2.4 | 3.47 | 2.23 | 271 | 242 |
| 13-07-23 | 2.47 | 3.67 | 2.37 | 25 | 2 |
| 13-07-23 | 3.83 | 5.6 | 3.23 | 744 | 542 |

Table S3: Spawn and size data for lab-grown *M. leidyi* from one pseudokreisel tank (Tank A)

| Date | Oral-Aboral Length (cm) | Overall Length (cm) | Pharyngeal Width (cm) | Number of Eggs | Number of Hatched Larvae |
|---|---|---|---|---|---|
| 17-07-23 | 3.23 | 5.2 | 3.1 | 659 | 250 |
| 17-07-23 | 1.8 | 2.53 | 1.87 | 63 | 6 |
| 17-07-23 | 2.3 | 3.83 | 2.5 | 553 | 497 |
| 17-07-23 | 1.7 | 2.47 | 1.67 | 78 | 49 |
| 18-07-23 | 2.8 | 4.23 | 2.63 | 626 | 547 |
| 18-07-23 | 1.87 | 2.87 | 1.97 | 75 | 21 |
| 18-07-23 | 1.5 | 2.23 | 1.87 | 2 | 0 |
| 18-07-23 | 3.37 | 5.63 | 3.1 | 453 | 316 |
| 19-07-23 | 2.43 | 3.67 | 2.43 | 239 | 181 |
| 19-07-23 | 3.2 | 4.87 | 2.9 | 1056 | 793 |
| 19-07-23 | 1.53 | 2.4 | 1.77 | 160 | 116 |
| 19-07-23 | 1.9 | 3.17 | 2.1 | 219 | 56 |
| 20-07-23 | 1.8 | 2.6 | 1.73 | 215 | 28 |
| 20-07-23 | 2.83 | 4.13 | 2.6 | 437 | 189 |
| 20-07-23 | 3.23 | 5.07 | 2.83 | 620 | 604 |
| 20-07-23 | 2.23 | 3.33 | 2.17 | 171 | 113 |

Table S4: Spawn and size data for lab-grown *M. leidyi* from one pseudokreisel tank (Tank B)
